# Frequency shapes the quality of tactile percepts evoked through electrical stimulation of the nerves

**DOI:** 10.1101/2020.08.24.263822

**Authors:** Emily L. Graczyk, Breanne P. Christie, Qinpu He, Dustin J. Tyler, Sliman J. Bensmaia

## Abstract

Touch is critical for our ability to manipulate objects, as evidenced by the deficits incurred when touch is absent. To restore the sense of touch via electrical stimulation of the peripheral nerves requires that we understand how the parameters of stimulation shape the evoked sensation. To this end, we investigated the sensory consequences of changing the frequency of pulse trains (PF) delivered to the peripheral nerves of humans chronically implanted with multi-channel nerve cuff electrodes. We found that increases in PF led to systematic increases in perceived frequency, up to about 50 Hz, at which point further changes in PF had little to no impact on sensory quality. Above this transition frequency, ratings of perceived frequency levelled off, the ability to discriminate changes in PF was abolished, and verbal descriptors selected to characterize the sensation changed abruptly. We conclude that the quality of electrically evoked tactile sensations can be shaped by imposing temporal patterns on a fixed neural population, but this temporal patterning can only be resolved up to about 50 Hz. These findings highlight the importance of spike timing in shaping the quality of a sensation and will contribute to the development of encoding strategies for conveying touch feedback through bionic hands and feet.

## Introduction

Physical interactions with our environment give rise to a rich and complex tactile experience. While a touch can be readily localized to a specific location on the body and described as light or strong, its sensory quality is multifaceted and often difficult to describe. For example, the tactile experience of a fabric is different from that of a vibration. Sensory quality does not fall on one continuum or even a few. Indeed, even a single sensory subspace, such as that for tactile texture, is complex and can be further broken down into component dimensions and subspaces^1–4^.

The neural underpinnings of sensory quality are also difficult to identify. The perceived location of a touch is determined by the receptive field location of the activated neural population^5–7^. The magnitude of a percept is determined by the overall population spike rate evoked in the peripheral nervous system^8,9^. Quality, on the other hand, is determined by the specific spatiotemporal pattern of activation elicited in the nervous system. Some patterns of activation give rise to a texture percept, others to a motion percept, and these patterns can co-occur and be multiplexed in the nerve^10^.

One determinant of quality is the degree to which each class of tactile nerve fibers is activated by the stimulus. Indeed, the sensation evoked when an individual nerve fiber is electrically activated depends on its afferent class – whether it is slowly-adapting or rapidly-adapting, type I or type II^5,11^. As a result, one reason different tactile stimuli evoke sensations of different qualities is because they differentially activate different classes of tactile afferent fibers. For example, skin vibrations recruit afferent populations in a frequency-dependent manner^12,13^. This differential recruitment accounts in part for the frequency-dependence of vibrotactile pitch, the qualitative dimension associated with changes in vibratory frequency^12–14^.

Another determinant of quality is the temporal patterning in the spiking response. For example, the perception of texture is shaped by the millisecond-level timing of sequences of spikes evoked in nerve fibers^15^. Similarly, phase-locked responses of nerve fibers to skin vibrations underlie their perceived vibrotactile pitch, which complements the information about frequency carried in the pattern of recruitment across classes^16,17^. In natural mechanical touch, the co-occurrence of these coding principles – differential recruitment and temporal patterning – makes it difficult to disentangle their respective roles in the determination of sensory quality.

Electrical stimulation offers a unique opportunity to examine the contribution of spike timing on tactile perception independently of the relative recruitment of afferent classes. An electrical pulse train delivered to the nerve will synchronously activate local neurons regardless of fiber class, as there are no known class-specific properties to enable preferential recruitment^18,19^. The impact of stimulation frequency on quality is thus mediated by the phase-locked spiking response, as all fiber classes are equally likely to be activated. Thus, by varying stimulation frequency in the peripheral nerve, we can directly examine the role of spike timing on quality perception.

Electrical stimulation of somatosensory peripheral nerves evokes vivid tactile sensations that are experienced on the hand^20–24^. While increasing stimulation frequency in the peripheral nerve is known to increase the perceived intensity of the percept, mediated by a concomitant increase in the population firing rate^9^, the effect on quality has not been previously studied. To fill this gap, we sought to understand how temporal patterns of activation in the nerve affect the quality of tactile percepts. We approached this question in two sets of psychophysical experiments with human amputees implanted with multi-channel nerve cuff electrodes. First, we investigated how changes in the frequency of electrical stimulation impact the perceived frequency of the stimulus. In these experiments, participants judged the evoked percept along a single dimension. We also assessed the sensitivity to changes in frequency. Second, we investigated the impact of stimulation pulse frequency on a multidimensional quality space defined by verbal descriptors to achieve a more holistic understanding of how pulse frequency shapes the quality of an electrically-evoked percept. Understanding how spike timing shapes quality will help unravel the peripheral neural code of touch and will lead to the development of approaches to convey a richer, more natural touch experience to individuals with bionic hands and feet.

## Results

Tactile sensations were elicited by delivering electrical pulse trains to the somatosensory nerves, either via Flat Interface Nerve Electrodes (FINEs) or Composite Flat Interface Nerve Electrodes (C-FINEs) (Figure 1a). Four unilateral upper limb amputees and two unilateral lower limb amputees participated in the study.

**Figure 1.**
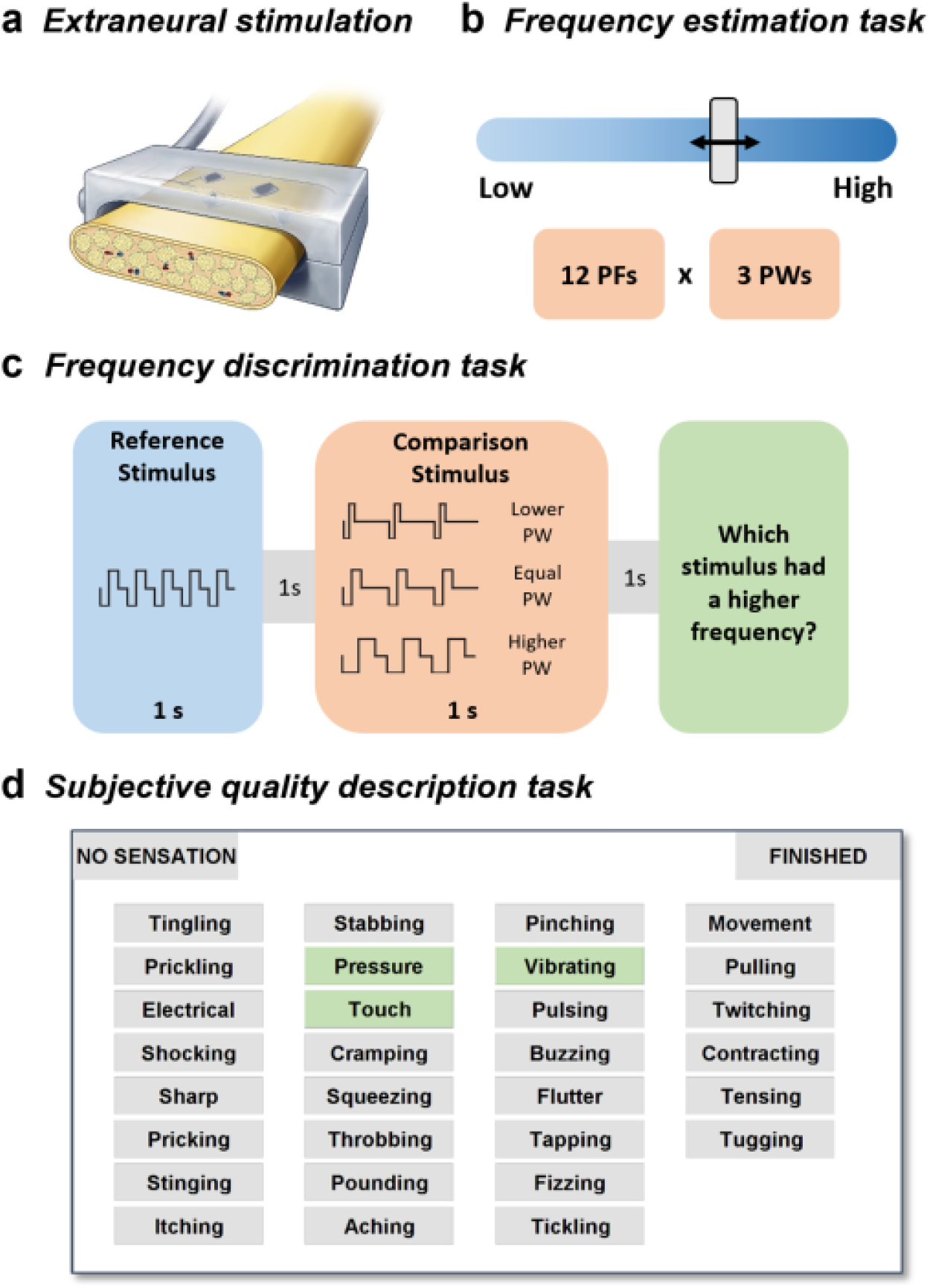
Experimental design. a) Electrical stimulation was delivered by an external neurostimulator through percutaneous leads to FINEs or C-FINEs implanted on the residual peripheral nerves of four trans-radial and two trans-tibial amputees. Stimulation consisted of trains of square, biphasic, charge-balanced pulses. b) Frequency estimation task. Three participants were asked to estimate the perceived frequency of an evoked percept by moving a slider along a horizontal bar. Stimuli varied in both stimulation pulse frequency (PF) and pulse width (PW). c) Frequency discrimination task. Four participants judged which of two sequentially-presented pulse trains was higher in perceived frequency, while ignoring any differences in perceived intensity. d) Quality description task. Six participants selected sets of words that described the quality of the evoked percept. Participants could select as many words as they wished to describe each stimulus.

### Perceived frequency increases with pulse frequency up to ∼50 Hz

First, we assessed the degree to which changes in frequency can be judged along a single perceptual continuum. To this end, we delivered stimuli that varied in pulse frequency (PF) and asked participants to rate the perceived frequency by positioning a slider along a horizontal bar (Figure 1b). To ensure that participants rated frequency rather than intensity, we varied the stimulation pulse width (PW), which also modulates perceived intensity^9^, from detection threshold to the maximum comfortable level. If frequency estimates were based solely on perceived intensity, then ratings would increase with increases in either PF or PW. PWs were scaled based on the detection threshold, which was measured for each contact separately.

At low PFs, perceived frequency increased logarithmically with PF and was independent of PW (Figure 2). At higher PFs, perceived frequency remained constant or even decreased with increases in PF and was instead modulated by PW (Figure 2b). We fit piecewise logarithmic functions to the relationships between perceived frequency and PF to determine the transition point between the rising and falling (or sustained) portions of the relationships (R^2^ = 0.85 ± 0.06, mean ± SEM) (Figure 2a). We found the transition frequency to be 60.0 ± 1.1 Hz (mean ± SEM). Transition frequency was highly consistent across participants and contacts and was not significantly modulated by PW (ANOVA, F(2,12) = 0.59, p = 0.57).

**Figure 2.**
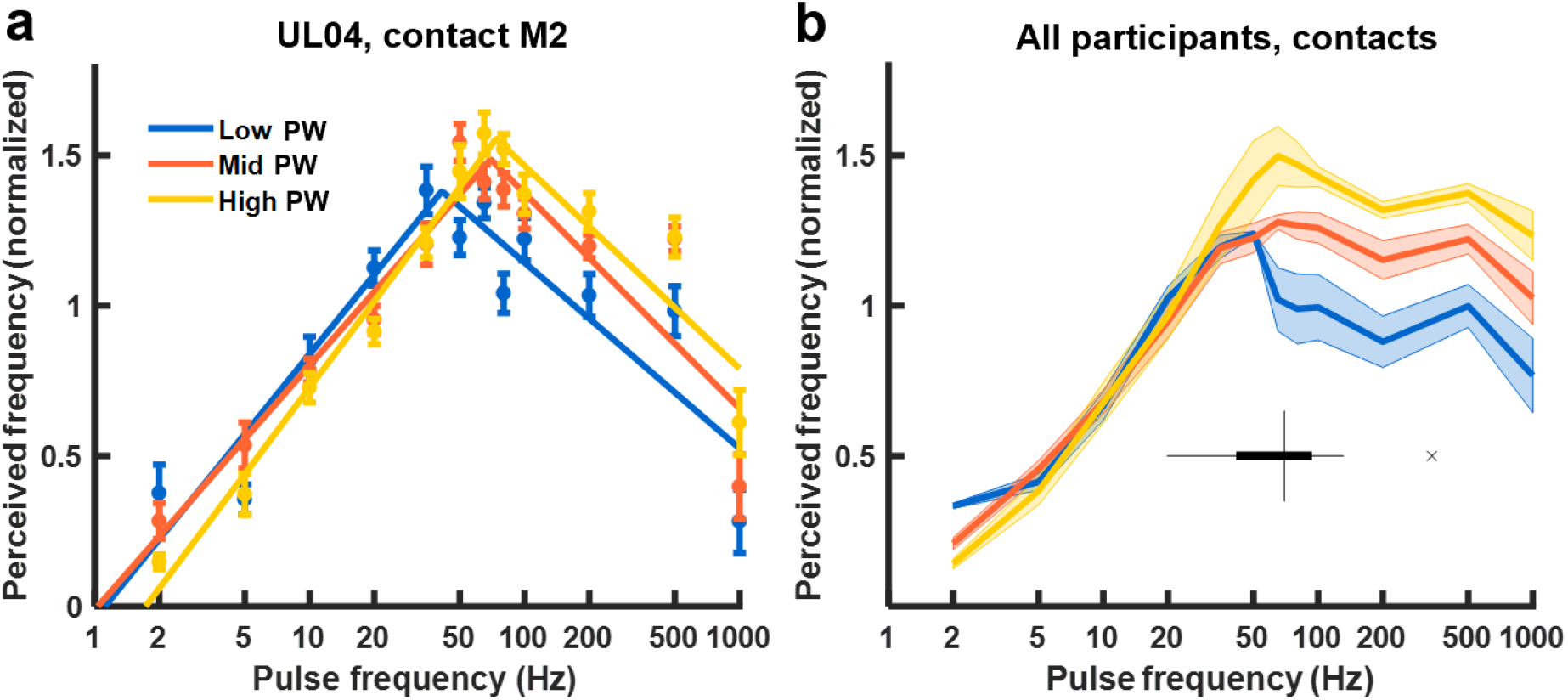
Perceived frequency estimates. a) Perceived frequency versus PF for a representative electrode contact (UL04, contact M2). Filled circles denote average perceived frequency ratings (n=15 per condition), normalized to the mean within experimental block (6 blocks). Color denotes the PW level and error bars denote the standard error of the mean. Solid lines represent piecewise logarithmic functions fit to the data. b) Average perceived frequency across all electrode contacts and participants versus stimulation PF (n=5). Color denotes the PW level and the shaded region corresponds to the standard error of the mean. The boxplot depicts the median and interquartile range of the transition frequency in the piecewise functions across participants and contacts (n=15). One outlier, denoted with an x, was removed from the analysis (see Methods). Note that the abscissa is logarithmic.

Having identified two different regimes in the PF response characteristic, we then examined the impact of PF and PW on perceived frequency in each regime separately. Below the transition PF, perceived frequency was significantly dependent on PF (F(5,82) = 27.83, p < 0.001) but not PW (F(2,82) = 1.58, p = 0.212). Above the transition PF, perceived frequency ratings depended on PW (F(2,82) = 14.08, p < 0.001) but not PF (F(5,82) = 1.33, p = 0.262). In summary, changes in PF exert a systematic effect on perceived frequency up to about 60 Hz, and this perceptual effect can be clearly distinguished from perceived magnitude. Increases in PF beyond 60 Hz do not change the quality of the percept along the perceived frequency continuum.

### Frequency discrimination is abolished above ∼50 Hz

Having found that the effects of PF on electrically evoked tactile percepts occupy two regimes – one at low and one at high frequencies – we next examined whether these regimes might be reflected in the participants’ ability to discriminate *changes* in PF. To this end, we asked participants to judge which of two sequentially presented pulse trains was higher in perceived frequency (Figure 1c). In each experimental block, which consisted of 270 trials, a standard stimulus (at 20, 50, or 100 Hz) was paired with a comparison stimulus whose PF and PW varied from trial to trial over a range (25-175% of the standard PF). The comparison PW took on one of three values: one shorter than, one equal to, and one longer than the standard PW (scaled based on the threshold PW, see Methods). The variation in PW was intended to reduce or abolish the informativeness of perceived magnitude, which is modulated by changes in both PW and PF^9^.

#### Discrimination with a constant pulse width

First, we examined participants’ ability to distinguish changes in PF independently of PW by only analyzing same-PW pairs. We found that participants could discriminate on the basis of frequency on most electrode contacts when the standard PF was set at 20 Hz or 50 Hz, as evidenced by smooth psychometric functions over the range tested (Figure 3a & b; see Supplementary Figure 1a, b for exemplary “poor” contacts). With the 100-Hz standard, however, PF discrimination performance was very poor on all contacts, and participants never achieved criterion performance (75% correct, Figure 3c, Supplementary Figure 1 d, e, f, see Methods). The just noticeable difference (JND) – defined as the difference in PF that yields 75% correct discrimination – was 3.867 ± 0.51 Hz and 9.250 ± 0.71 Hz (mean ± SEM) for the 20- and 50-Hz standards, respectively, yielding nearly constant and statistically indistinguishable Weber fractions (0.193 ± 0.03 Hz and 0.185 ± 0.01 Hz) (Figure 3e). In contrast, JNDs were undefined for the 100-Hz standard for all tested contacts. Note that participants could perceive the stimuli even when the JND was undefined but they could not discriminate between them.

**Figure 3.**
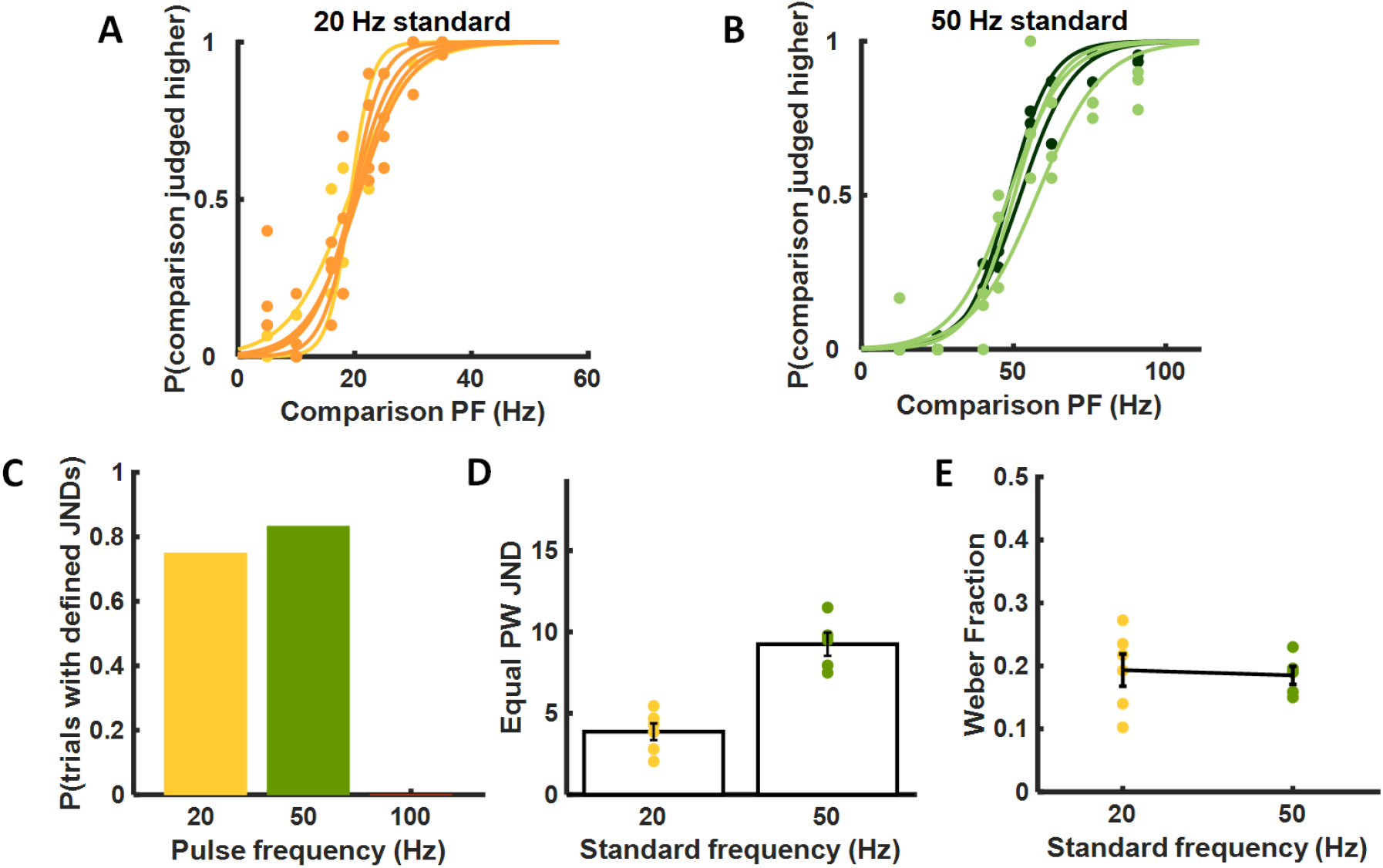
PF discrimination performance with constant PW. a) Performance with the 20-Hz standard for participants UL02 (yellow) and UL04 (orange). Each curve denotes the mean performance with a different electrode contact (n=6). b) Performance with the 50-Hz standard across contacts (n=5) for UL02 (light green) and UL04 (dark green). c) Proportion of stimulation contacts yielding defined JNDs for the three standard frequencies when comparison and standard PWs are equal. Participants achieved threshold performance on 75% (6 out of 8) contacts tested with the 20-Hz standard and 83% (5 out of 6) contacts tested with the 50-Hz standard. Participants never achieved criterion performance (0 out of 7) with the 100-Hz standard. d) Just noticeable differences (JNDs) for the equal PW conditions with the 20-Hz (n=6) and 50-Hz (n=5) standards. e) The Weber fractions for the 20-Hz (n=6) and 50-Hz (n=5) standards were statistically indistinguishable.

#### Effect of pulse width on frequency discrimination

Second, we examined the degree to which PF could be discriminated independently of changes in perceived magnitude. Trials with equal-PW pairs were interleaved with trials in which the PW of the comparison was different from that of the standard stimulus. To the extent that participants relied on differences in perceived magnitude to make their frequency judgments, PW would systematically bias participants’ frequency judgments, thereby causing lateral shifts in psychometric functions. Changes in PW may also lead to changes in the clarity or vividness of the evoked percepts, thereby resulting in changes in the discriminability of PF.

We observed systematic leftward shifts in the psychometric functions when the comparison PW was lower than the standard PW and systematic rightward shifts when the comparison PW was higher than the standard PW (Figure 4a, b, d & e). In other words, pulse trains with lower PW tended to be perceived as higher in frequency across PFs. This bias was quantified by computing the point of subjective equality (PSE) – the PF at which participants were equally likely to select the standard as they were the comparison stimulus. The PSE increased as the comparison PW increased for both the 20-Hz and 50-Hz standards (Figure b & e, repeated measures ANOVA, F(2,16) = 17.34, p < 0.001; F(2,14) = 17.46, p = 0.0012, respectively). In summary, the participants exhibited a slight tendency to select the more intense stimulus as being lower in frequency, consistent with previous findings that the pitch of a vibrotactile stimulus decreases as the stimulus amplitude increases^25^. Note that this tendency was also observed in the frequency estimation ratings but was too weak to achieve statistical significance.

**Figure 4.**
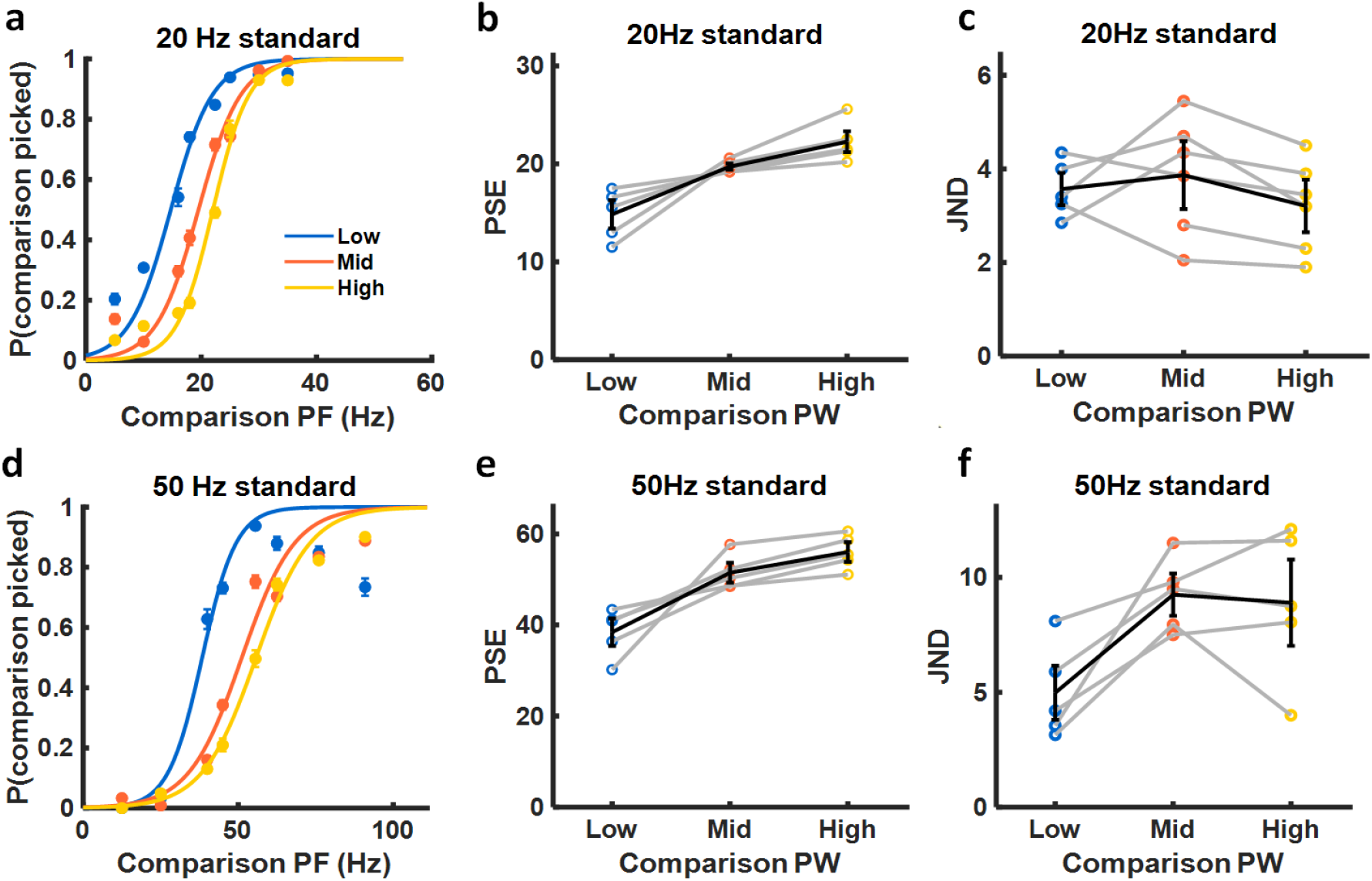
PF discrimination performance with variable PW. a) Performance with the 20-Hz standard and three comparison PWs, averaged across electrode contacts (n=6). b) Point of subjective equality (PSE) vs. PW for the 20-Hz standard (n=17). Gray lines denote different contacts, black line denotes the mean across contacts and participants. PSE increases with PW, revealing a PW-dependent bias in PF discrimination performance. Participants exhibited a tendency to perceive higher PWs as being lower in PF. c) JND vs. PW for the 20-Hz standard (n=17). There was no significant effect of PW on JND with the 20-Hz standard. d) Performance with the 50-Hz standard and three comparison PWs, averaged across electrode contacts (n=5). e) Point of subjective equality (PSE) vs. PW for the 50-Hz standard (n=15). PSE was significantly higher at higher PWs. f) JND vs. PW for the 50-Hz standard (n=15). JNDs tended to be higher at higher PWs.

Next, we investigated the degree to which the comparison PW affected participants’ sensitivity to changes in PF by examining the effect of PW on the JND. For the 20-Hz standard, we found no systematic effect of PW on JND (Figure 4c, repeated measures ANOVA, F(2,16) = 1.35, p = 0.3082). For the 50-Hz standard, on the other hand, JNDs increased as PW increased (Figure 4f, F(2,14) = 9.58, p = 0.0075). In other words, participants became somewhat less sensitive to changes in PF at higher PWs, but only for the higher-PF standard.

### Subjective reports of sensory quality depend on stimulation frequency

Given that PF reliably influenced perceived quality along a single frequency continuum, we then investigated how PF shapes subjective quality more broadly. To this end, participants indicated which subset of a list of 30 qualitative descriptors pertained to each stimulus. Participants were encouraged to select as many words as necessary to describe the electrically evoked sensation.

#### Qualitative reports

On average, participants selected 6.5 ± 2.6 words (mean ± SEM) to describe each sensation. The total number of words chosen to describe a stimulus varied across participants, ranging from 5 to 22.

Examination of the participants’ word selections revealed that stimulation PF influenced the descriptors chosen to characterize the stimuli. That is, different sets of words were selected to describe low-frequency stimuli than high-frequency stimuli (Figure 5, Supplementary Table 1), as has been found with vibrotactile stimulation^26^ and intracortical microstimulation^27^. For example, descriptors related to periodic sensations, such as ‘tapping’, ‘pulsing,’ ‘twitching,’ and ‘flutter,’ were commonly selected for low-PF stimuli, whereas continuous sensations, such as ‘tingling,’ ‘buzzing,’ ‘electrical,’ ‘pressure,’ and ‘touch,’ were reported for high-PF stimuli (Supplementary Table 1). In addition, the total number of reported descriptors increased as a function of stimulation PF (linear regression, p<0.001).

**Figure 5.**
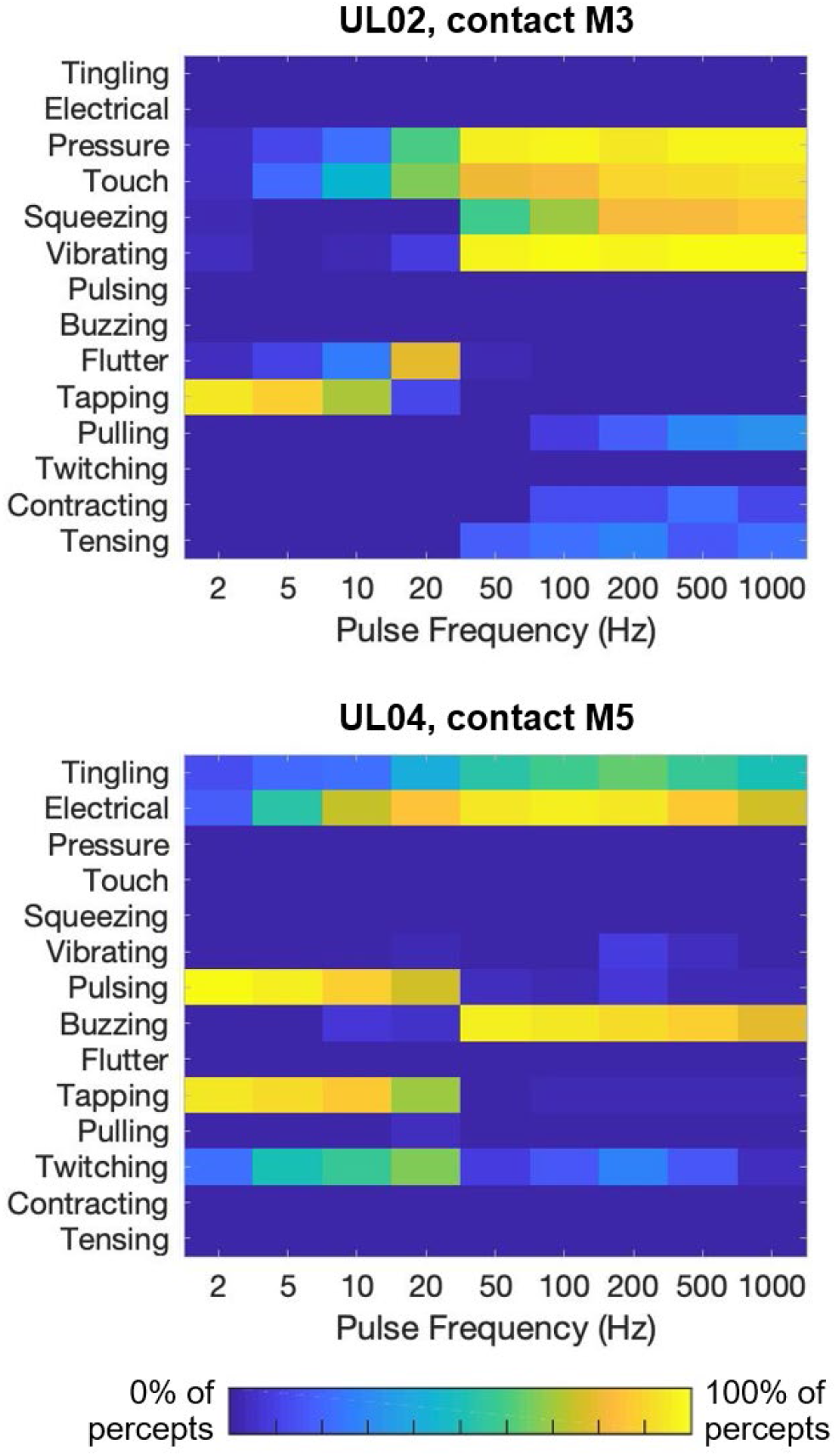
Quality descriptors reported for electrically-evoked percepts at each PF, averaged across PWs. Color denotes the proportion of times a descriptor was selected at each PF (top: contact M3/UL02; bottom: contact M5/UL04). Participants used different words to describe their sensory experiences. However, descriptors were similar across contacts for each participant (see Supplementary Figure 2).

Participants used similar descriptors across electrode contacts, as evidenced by highly correlated descriptor selection within-participants (R^2^ = 0.66 ± 0.08, mean ± SEM). The consistency in reports across electrode contacts indicates that quality is largely independent of which region of the nerve is stimulated. However, the descriptors ascribed to individual stimuli varied widely from participant to participant (Figure 5, Supplementary Figure 2), yielding weak correlations in descriptor selections across participants (R^2^ = 0.14 ± 0.04). These variations likely reflect the participants’ idiosyncratic descriptor preferences, since some words, such as tapping and pulsing, may be construed as synonymous.

#### A robust representation of sensory quality

To overcome participant-specific descriptor choice and assess the effect of PF on quality across participants, we computed differences in descriptor selection between each pair of stimuli. To this end, we first represented quality as a 30-element vector, where each element corresponded to the proportion of trials in which a given descriptor was selected. We then ran a principal component analysis (PCA) on the vectors of individual participants to remove redundant or highly correlated descriptors. Finally, we computed the Euclidean distance between each pair of stimulation conditions (PW and PF combinations) in this reduced dimensional space. These distances represent the perceived dissimilarity between pairs of stimuli, estimated from the differences in descriptor selection (Figure 6a,b).

**Figure 6.**
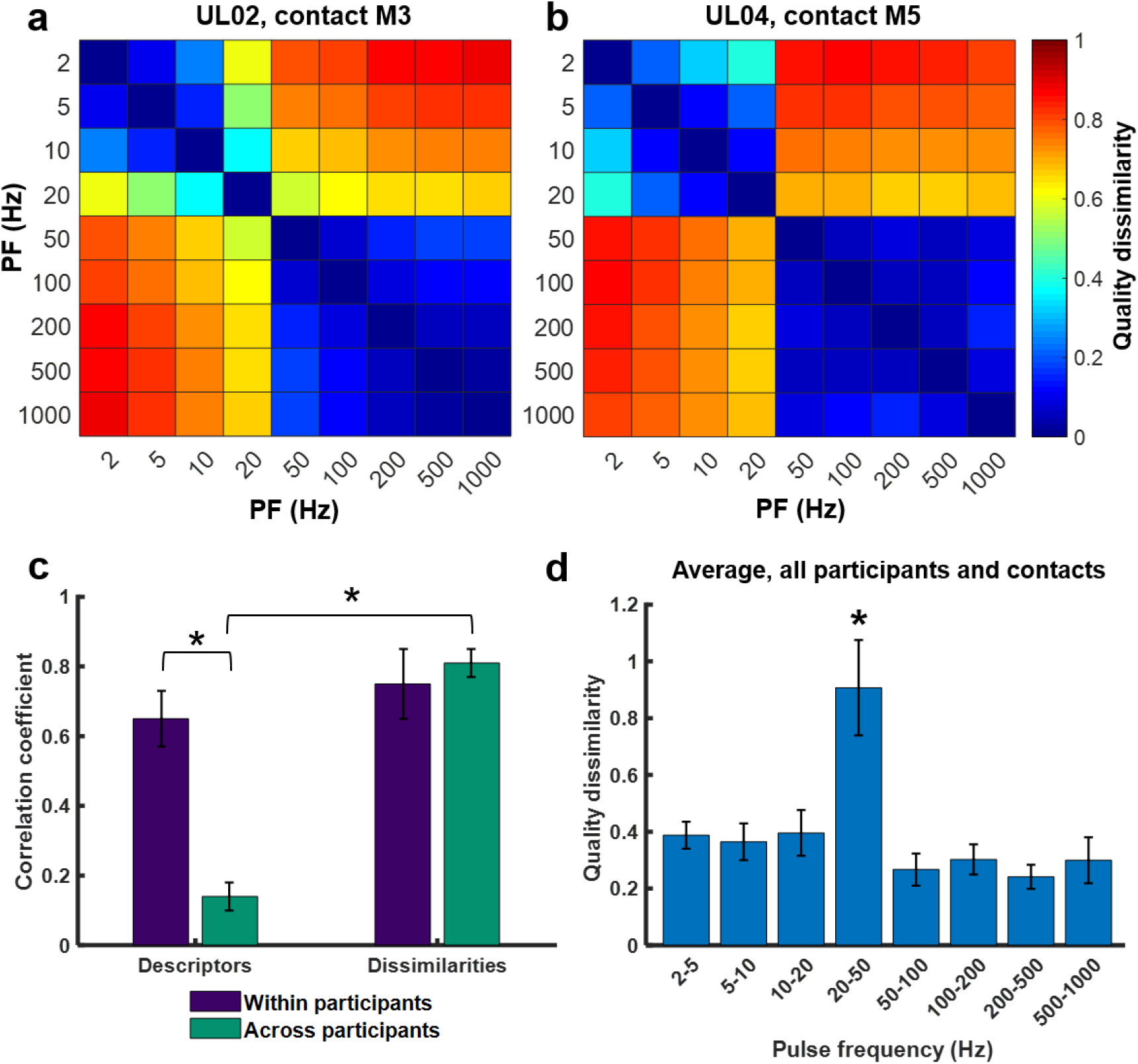
Perceived dissimilarity estimated from differences in descriptor usage. a) Perceived dissimilarity of stimuli varying in PF for contact M3 of participant UL02. b) Perceived dissimilarity of stimuli varying in PF for contact M5 of participant UL04. Note that the contacts/participants shown in a and b are the same as in Figure 5. c) Within-participant (n=8, purple) and across-participant (n=15, teal) pairwise correlations for two metrics of quality. “Descriptors” denotes the words selected by the participant for each stimulus, and “Dissimilarities” denotes the Euclidean distance between pairs of stimulation conditions. Error bars denote the standard error of the mean. Asterisks denote statistically significant differences with p<0.001. d) Perceived dissimilarity between successive PF conditions, averaged across PWs (n=33). Error bars indicate the standard error of the mean. The asterisk denotes significance at p<0.001.

We found that the resulting dissimilarity matrices were highly consistent across participants and electrode contacts (Figure 6a,b). Indeed, dissimilarity matrices were significantly more similar across participants (R^2^ = 0.81 ± 0.04) than were the descriptor selections from which they were derived (R^2^ = 0.14 ± 0.04) (One-way ANOVA, F(3,42) = 35.80, p < 0.001) (Figure 6c, teal bars). The consistency in dissimilarity matrices across participants suggests that the relationship between sensation quality and PF is similar across participants, despite idiosyncratic descriptor usage.

#### Impact of PF on overall quality

Having established a robust representation of multi-dimensional quality based on dissimilarity, we then assessed the degree to which changes in PF led to changes in the quality of the evoked percept. We found a sharp transition in sensation quality between 20- and 50-Hz for all tested contacts. In fact, the dissimilarity between 20-Hz and 50-Hz stimuli was significantly greater than the dissimilarity between any other pair of adjacent frequencies (ANOVA, F(7,197) = 16.14, p < 0.001) (Figure 6d). In other words, the perceptual quality was consistent across PFs ranging from 2 to 20 Hz and across PFs from 50 to 1000 Hz, but these two subsets of stimuli felt very different from one another. There was no systematic effect of stimulation PW on the dissimilarities between successive PFs (ANOVA, F(2,197) = 1.22, p = 0.3).

We then determined the extent to which differences in PF and PW resulted in differences in the evoked sensation. To quantify the impact of PF and PW on quality, we averaged the dissimilarity matrices across contacts and participants (Figure 7a, Supplementary Figure 3) and regressed the resulting matrix on differences in PF or PW (ΔPF, ΔPW). The regression model that included both ΔPF and ΔPW yielded accurate predictions of dissimilarity (R^2^ = 0.67, F(3,725) = 499, p < 0.001) (Figure 7b). While both ΔPF and ΔPW had significant impacts on dissimilarity, the contribution of ΔPF was much higher than that of ΔPW (standardized coefficients: β_ΔPF_ = 0.17, p < 0.001; β_ΔPW_ = 0.05, p < 0.001) (Figure 7c). The interaction between ΔPF and ΔPW was significant but weak (β_ΔPF*ΔPW_ = -0.04, p < 0.001).

**Figure 7.**
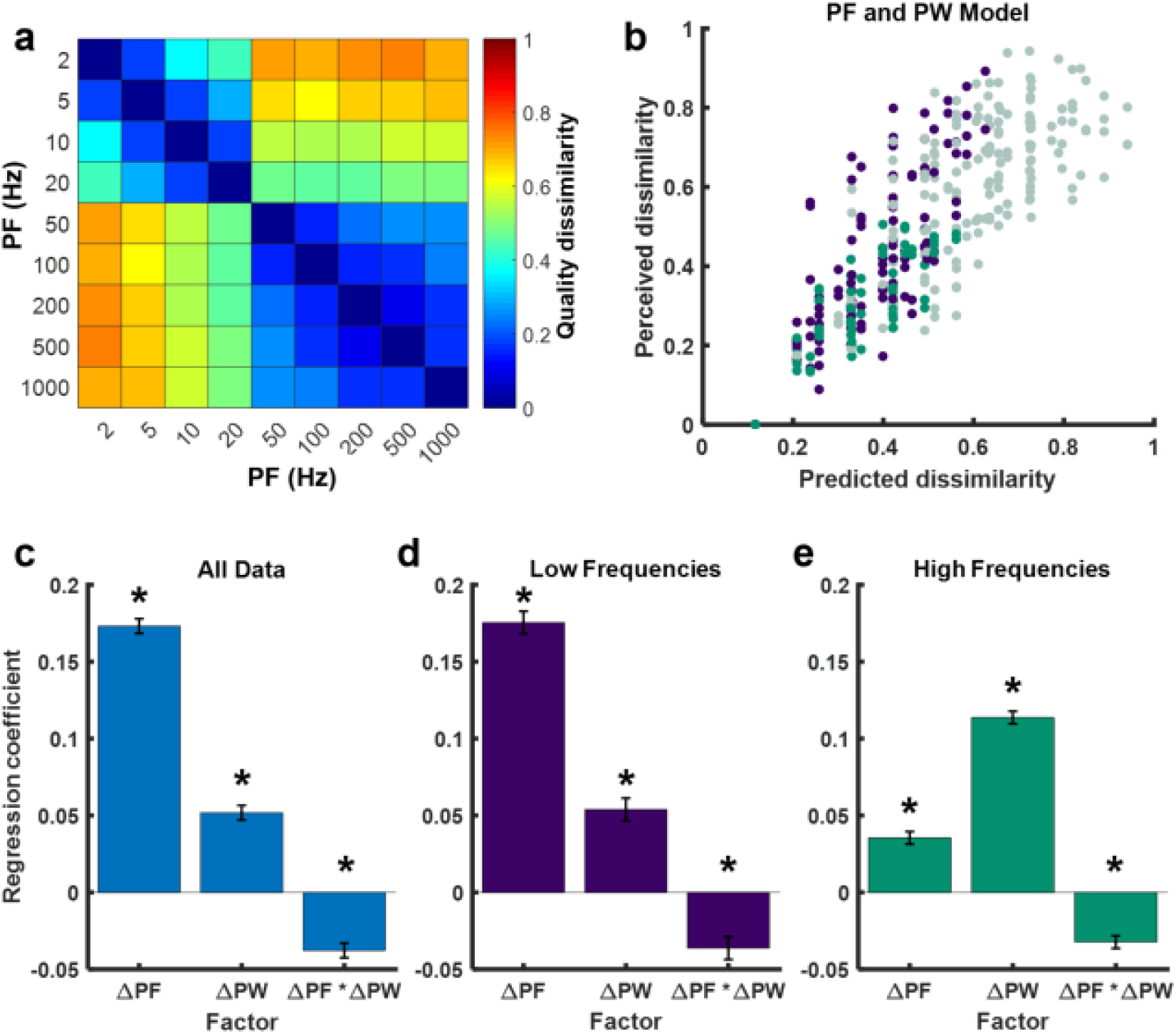
Contribution of stimulation parameters to sensation quality. a) Dissimilarity matrix averaged across participants and contacts (n=11). b) Measured dissimilarity vs. prediction from a linear combination of ΔPF, ΔPW, and their interaction. Purple points represent dissimilarities between stimuli at low frequencies (<50 Hz), green points represent dissimilarities between stimuli at high frequencies (>50 Hz), and grey points represent dissimilarities between high and low frequency stimuli. c) Standardized regression coefficients for the model with all data included. d) Regression coefficients for the model with only low frequency data included (<50 Hz). e) Regression coefficients for the model with only high frequency data included (>50 Hz). c,d,e) Error bars are standard error of the estimates. Asterisks denote significant contribution to the model at p<0.001.

Given the sharp transition in perceived quality around 50 Hz, we fit separate regression models for the dissimilarity data below 50 Hz and above 50 Hz. At low PFs, the regression was nearly identical to that obtained using the full dataset (R^2^ = 0.74, F(3,221) = 214, p < 0.001; β_ΔPF_ = 0.18, p < 0.001; β_ΔPW_ = 0.05, p < 0.001; β_ΔPF*ΔPW_ = -0.04, p < 0.001) (Figure 7d). In contrast, at high PFs, the effect of PW dominated that of PF (R^2^ = 0.74, F(3,140) = 302, p < 0.001; β_ΔPF_ = 0.04, p < 0.001; β_ΔPW_ = 0.11, p < 0.001; β_ΔPF*ΔPW_ = -0.03, p < 0.001) (Figure 7e). We performed the same analyses on individual participants’ data and obtained the same results, confirming that these results were not an artifact of pooling (Supplementary Figure 4).

#### Relationship between perceived frequency and dissimilarity

Finally, we determined whether the effect of PF on dissimilarity could be explained solely based on its effect on perceived frequency. That is, we examined whether PF modulates perceptual quality along a single continuum. To this end, we first regressed dissimilarity on the perceived frequency ratings obtained in the first experiment, then assessed whether the residuals co-varied with ΔPF. In the low-frequency regime, we found that differences in perceived frequency predicted dissimilarity ratings well (R^2^ = 0.71, F(1,223) = 545, p < 0.001) (Supplementary Figure 5a) and that ΔPF was not significantly predictive of the residuals (R^2^ = -0.004, F(1,223) = 0.07, p = 0.80) (Supplementary Figure 5c). Thus, the effect of PF on quality is confined to a single continuum in the low frequency regime.

In the high-frequency regime, we similarly found that perceived frequency was a significant predictor of dissimilarity, as it was in the low frequency regime (R^2^ = 0.53, F(1,142) = 164; p < 0.001) (Supplementary Figure 5b). However, ΔPF continued to have a weak but significant effect on dissimilarity ratings after regressing out the effect of perceived frequency (R^2^ = 0.07, F(1,142) = 11, p = 0.001) (Supplementary Figure 5d). Thus, in the high-frequency regime, the effect of PF on quality dissimilarity cannot be explained purely on the basis of perceived frequency, consistent with the finding that perceived frequency is not dependent on PF at those frequencies (see Figure 2). Note, however, that the overall impact of PF on quality was weak in the high-frequency range (see Figure 7d), so the magnitude of these effects was small.

## Discussion

### Stimulation frequency shapes sensory quality below 50 Hz

We find that the temporal pattering in the neuronal response, which in this case was periodic and defined by PF, shapes the perceived quality of electrically-evoked tactile percepts both along a single continuum – perceived frequency – and across a multi-dimensional space defined by perceptual descriptors. The effect of PF on quality was most pronounced at low PFs, below approximately 50 Hz. In this frequency range, increases in PF led to increases in perceived frequency and systematic changes in verbal reports of sensory quality. In addition, we found that the impact of PF on verbally reported quality could be accounted for by variations in perceived frequency, demonstrating that PF modulates quality along a single continuum. The fact that PW did not have a significant effect on perceived frequency and had a marginal effect on overall quality in this frequency range indicates that the effect of PF on quality does not simply reflect a change in perceived magnitude.

Taken together, these results show that temporal patterning in the aggregate afferent response can modulate the quality of tactile percepts at low frequencies, despite the fact that this temporal patterning is distributed across afferents of all classes. Thus, the differential recruitment of afferent classes based on their frequency sensitivity profiles is not the only factor that shapes quality perception, though it likely plays an important role^5,28^.

### Changes in PF are indiscriminable at high frequencies

Changing the frequency of vibrations delivered to the skin results in a change in the sensory experience along a continuum described as ‘vibrotactile pitch,’ underscoring its similarity to the sensory consequences of changes in the frequency of an acoustic tone^25,29^. Human observers can reliably discriminate a change in vibrotactile frequency of around 20%, but sensitivity to changes in frequency decreases at higher vibration frequencies^30–33^. Changes in vibrotactile frequency also produce changes in perceived magnitude^34^. However, human participants can discriminate vibratory frequency in the presence of concomitant and independent changes in vibratory amplitude^35^, demonstrating that vibrotactile frequency also shapes the quality of the percept.

The ability to perceive the frequency of a vibration is thought to be mediated by a timing code in the nerve. Sinusoidal vibrations delivered to the skin elicit periodic, phase-locked spiking responses in tactile nerve fibers, and the frequency of these responses matches the frequency of stimulation^16,26^. When this temporal patterning is absent, for example at low stimulus amplitudes, the ability to discriminate frequency is abolished^36^.

Pulsed electrical stimulation of the nerve also produces a highly periodic response in the activated tactile nerve fibers^37^ and this frequency-dependent temporal patterning likely mediates the ability to discriminate the perceived frequency of electrical pulse trains. In fact, sensitivity to changes in the frequency of electrical stimulation is similar to its vibrotactile counterpart, both yielding Weber fractions of ∼0.2. We also demonstrate that stimulation PF has a larger effect on the quality of an electrically-evoked sensation than it does on its magnitude: the Weber fraction for PF along the perceived magnitude continuum (0.3)^9^ is higher than that along the perceived frequency continuum.

In contrast to vibrotactile frequency discrimination, which can be achieved over a wide range of frequencies up to 400 Hz and beyond^14,35,38^, we show that frequency discrimination of peripheral nerve stimulation is only possible at frequencies below ∼50 Hz. Above this point, frequency discrimination falls to near chance. This is especially interesting in light of the fact that non-human primates can distinguish the PF of intracortical microstimulation up to around 200 Hz, though sensitivity to PF changes decreases at high frequencies^39^. In addition, while our results showed that pulse trains with higher PW tended to be judged as lower in frequency, prior studies of intracortical microstimulation in non-human primates found that stimuli with higher pulse amplitude (PA) tended to be reported as being higher in frequency^38^. Assuming that PW and PA have similar neural and sensory correlates, the impact of intensity on frequency discrimination is thus different for the nerve and cortex. The basis for these differences is unclear, but the neural consequences of stimulation differ between nerve – a bundle of independent channels – and cortex – a densely interconnected circuit.

### Perceived dissimilarity is a robust way to quantify quality

Quality is difficult to quantify because this aspect of a sensory experience occupies a high dimensional space and measuring it relies on verbal reports, which are often highly idiosyncratic. In this study, the specific words chosen to describe the qualitative sensory experiences varied widely across participants. To mitigate this variability in descriptor selection, we developed an approach to represent quality based on *differences* in the descriptors that participants ascribed to the sensations. We found that this representation of quality was very similar across participants, suggesting that tactile sensations evoked from neural stimulation may be approximately equivalent across individuals despite idiosyncratic descriptor usage. We propose that our approach could be used in future studies of quality perception to improve comparisons across participants when the perceptual space is not confined to one or a few experimenter-defined dimensions.

### Quality abruptly changes at 50 Hz

As discussed above, the ability to discriminate PF was abolished beyond about 50 Hz, and the influence of PF on quality was far more pronounced at low PFs. In contrast, PW had a greater impact on quality at high PFs than did PF. Furthermore, multi-dimensional quality also shifted abruptly at ∼50 Hz.

There are several possible explanations for the transition between the low and high frequency regimes (see Supplementary Notes), but the most likely possibility is that the unnatural activation of the afferent submodalities by peripheral nerve stimulation obscures the temporal pattern imposed by stimulation at high PFs. Electrical stimulation recruits Aβ fibers based on their fiber diameter and nodal voltages^40,41^, regardless of their submodality class. In natural touch, the various submodalities of tactile fibers differ in the frequency profiles of their sensitivity to skin vibrations: slowly adapting type 1 (SA1) fibers respond preferentially to low-frequency vibrations, Pacinian corpuscle-associated (PC) fibers respond preferentially to high-frequency vibrations, and rapidly adapting fibers (RA) fibers exhibit intermediate frequency preferences ^13^. A high-PF pulse train will thus activate SA1 fibers, and perhaps RA fibers, to a much greater extent than would a high-frequency vibration delivered to the skin. This large, unnatural signal may obscure the critical signal carried by appropriately responding afferents, especially PC fibers. This phenomenon would be further magnified by the fact that RA1 and SA1 fibers far outnumber PC fibers in the periphery (a ratio of 1:1.6:0.6 for SA1:RA1:PC fibers)^42^.

The hypothesis that over-activation of SA1 and RA fibers could be implicated in poor frequency perception at high PFs is supported by previous studies of intraneural microstimulation: while increases in the PF delivered to individual RA afferents induces increases in perceived frequency at low PFs, further increases only lead to increased intensity^14^. Stimulation of individual SA1 fibers leads to sustained percepts at all but the lowest PFs^14^. The submodality over-activation hypothesis is also bolstered by the previous finding that the impact of peripheral afferent signals on cortical neural activity is highly dependent on the submodality of the peripheral afferents: PC signals in the periphery drive the *temporal patterning* of neural responses in cortex, whereas RA and SA1 signals instead drive the *strength* of the response in cortex^43^. Thus, the unnatural strength of the SA1 signal (and perhaps the RA signal) evoked via peripheral nerve stimulation may muddle the crucial frequency signal transmitted by PCs to the cortex. This phenomenon would be exacerbated at high PFs, at which SA1 fibers are typically relatively quiescent, consistent with our results. A similar phenomenon, in which RA signals interfere with SA1 signals about local geometric features, has been previously documented^44^.

### Dependence of quality on PF is consistent across fascicles

We did not observe any electrode-specific differences in the impact of PF on perceived quality, and prior studies similarly found no electrode-specific differences in intensity perception^9^. In contrast, the perceptual correlates of changes in PF of intracortical microstimulation have been shown to vary across electrodes in experiments with human subjects^27^ and non-human primates^38^. These results have been interpreted as reflecting somatotopic differences in the functions of the stimulated cortical circuits^28,45–47^. Conversely, the fascicles of the peripheral nerve display only minimal submodality-specific organization^48–50^, so the submodality composition of the nerve fibers activated through electrical stimulation is likely to be consistent across electrode contacts.

### Stimulation frequency can be employed to sculpt artificial touch for bionic limbs

A major challenge facing recent efforts to restore the sense of touch through neural stimulation is to evoke sensations endowed with an appropriate quality. The goal is for contact with a textured surface through a bionic hand to lead to a perceived sensation of the appropriate texture and contact with an edge to lead to the sensation of an edge. In the intact sensory system, the spatiotemporal pattern of activation evoked in the nerve when interacting with an object depends on the dynamics of contact and the properties of the object, and the evoked percept depends critically on these spatiotemporal neural patterns. While electrical stimulation of the nerves can reproduce the temporal aspects of this normal neural activity by patterning the pulses, it cannot mimic the complex spatiotemporal patterns due to limitations in the ability to selectively activate individual nerve fibers. Nonetheless, mimicking natural patterns of activation with biomimetic stimulation confers greater speed to certain sensory discrimination tasks performed with a prosthesis^51^ and improves performance on tasks involving object manipulation^52^. However, the impact of biomimetic stimulation on the quality of the evoked sensation is inconsistent^51–53^. Our results demonstrate that modulating stimulation frequency reliably and systematically modulates sensation quality. This phenomenon can be leveraged to convey information about object and substrate interactions more intuitively for upper and lower extremity neuroprostheses.

## Methods

### Participants

Six people volunteered for this study. Four participants had unilateral acquired amputations of the upper limb below the elbow and two had unilateral acquired amputations of the lower limb below the knee. All participants were implanted with multi-contact peripheral nerve cuff electrodes, either Flat Interface Nerve Electrodes (FINEs) or Composite Flat Interface Nerve Electrodes (C-FINEs)^54^. Upper limb participants (referred to as UL01-UL04) were implanted with 8-channel FINEs or 16-channel C-FINEs around the median, ulnar, and/or radial nerves between 3 and 7 years prior to participation in the present study. Contacts were selected for testing that elicited comfortable sensations on the anterior surface of the hand. Lower limb participants (LL01 and LL02) were implanted with 16-channel C-FINEs around the sciatic and tibial nerves between 1 and 3 years prior to participation in the present study. Contacts were selected for testing that elicited comfortable sensations on the plantar surface of the foot or posterior aspect of the ankle. Percutaneous leads connected the cuff electrodes to an external neurostimulator. Stimulation waveforms were square, biphasic, cathodic-first, and charge-balanced. Stimulation parameters were set in MATLAB (MathWorks, Inc.; Natick, MA, USA) and then sent to a computer running xPC Target (MathWorks, Inc.; Natick, MA, USA), which then drove the stimulator. The stimulator could produce stimulation with PW in the range of 5 to 255 µs in increments of 1 µs, PA in the range of 0 to 2 mA in increments of 0.1 mA, and inter-pulse intervals in the range of 1 to 10,000 ms in increments of 1 ms.

The Louis Stokes Cleveland Department of Veterans Affairs Medical Center Institutional Review Board and Department of the Navy Human Research Protection Program approved all procedures. This study was conducted under an Investigational Device Exemption obtained from the United States Food and Drug Administration. All participants provided written informed consent to participate in this study, which was designed in accordance with relevant guidelines and regulations.

### Frequency estimation task

In each trial, a 1-second pulse train was delivered and participants estimated the perceived frequency of the evoked sensation. Estimations were made using a visual analog scale displayed on a computer monitor, which displayed “Slowest possible frequency” on the far left of the scale and “Fastest possible frequency” on the far right of the scale. In the first trial, we advised participants to rank the perceived frequency somewhere in the middle of the scale. In subsequent trials, they were asked to rank perceived frequency as faster or slower with respect to prior stimuli. Participants were asked to ignore changes in intensity and location when making frequency estimates. A 3-second inter-trial interval was enforced to minimize the effects of perceptual adaptation^55^.

For each electrode contact, we tested twelve pulse frequencies (PF) and three pulse widths (PW). PFs were set to 2, 5, 10, 20, 35, 50, 65, 80, 100, 200, 500, and 1000 Hz. The three PWs were chosen on a per-contact basis to span the full range of suprathreshold, comfortable intensities. Briefly, we used a staircase procedure to find the minimum PA value that produced a detectable percept at the maximum PW of the stimulator (255 µs). We then used a two-alternative forced-choice tracking paradigm to find the minimum PW value that elicited a detectable percept at this threshold PA^9^. The “Low” PW value in the estimation task was the minimum PW that was reliably perceived at 2 Hz. The “High” PW value was chosen as the maximum PW that was comfortable at 500 Hz. The “Mid” PW value was chosen as the midpoint between the “Low” and “High” values. Preliminary tests with each contact indicated that these PWs resulted in clear differences in perceived intensity. The 36 stimuli were presented in pseudo-random order fifteen times per contact, split into six experimental blocks of 90 trials, all performed on the same day. Five electrode contacts across three participants were included in this analysis.

Frequency ratings were normalized by dividing each estimate by the within-block mean rating. Normalized ratings were then averaged within-contact for each PW and PF condition. We then fit piecewise linear functions to the base-10 logarithm of frequency, separated by PW condition. For each fit, coefficient estimates over their relevant ranges were randomly generated ten times, and the combination with the lowest mean squared error was selected. The transition between the rising and falling phases of the perceived frequency ratings was obtained from the optimized parameters. The transition metric was averaged across participants, contacts, and PWs after first removing an outlier that was more than two standard deviations above the mean. Means and standard errors of the frequency estimates were then calculated across participants and contacts within each PW and PF condition.

### Frequency discrimination task

Each trial consisted of two successive 1-second pulse trains separated by a 1-sec inter-stimulus interval. Participants judged which of the two sequentially presented pulse trains was higher in perceived frequency. In each experimental block, a standard stimulus (at 20, 50, or 100 Hz) was paired with a comparison stimulus whose PF and PW varied from trial to trial. The comparison PFs ranged from 25 to 175% of the standard PF. Specifically, for the 20-Hz reference, the comparison PFs were 5, 10, 16, 18, 20, 22.4, 25, 30, and 35 Hz; for the 50-Hz reference, the comparison PFs were 12.5, 25, 40, 45, 50, 55.55, 62.5, 76, and 90.9 Hz; and for the 100-Hz reference, the comparison PFs were 25, 50, 83, 90, 100, 111, 125, 145, and 166 Hz. The comparison PW took on one of three values: one shorter than, one equal to, and one longer than the standard PW. PWs were based on the same criteria as in the frequency estimation task. All PW values were between 70 to 130% of the standard PW. The standard stimulus was always at the intermediate PW. In each block, each stimulus pair was presented 10 times, and both the order of stimuli within the pair and the order of the pairs varied pseudo-randomly. The inter-trial interval was enforced to be at least 3 seconds long. Thus, each experimental block evaluated one electrode contact and consisted of 270 trials (9 comparison PFs × 3 comparison PWs × 10 repetitions). Eight contacts across four participants, six contacts across two participants, and six contacts across two participants were evaluated for the 20, 50, and 100 Hz standard, respectively.

Psychophysical performance was calculated as the proportion of trials in which the comparison stimulus was judged to be higher in perceived frequency than the standard stimulus. Psychometric curves were then constructed by fitting a cumulative normal density function to these proportions. Separate psychometric functions were fit for each standard frequency (20, 50, and 100 Hz). Just noticeable differences (JNDs) were calculated as the change in frequency required to achieve 75% correct performance. Two JND estimates were obtained from each psychometric function, one above and one below the standard, and these two estimates were averaged. Sessions in which performance did not achieve the 75% threshold performance criterion (above or below the standard) were considered to have an undefined or incalculable JND and were considered to be “poor” performance contacts. These data points were omitted from further analysis in the main results section and were only included in the supplementary materials. The point of subjective equality (PSE) was defined as the comparison frequency that corresponded to 50% correct performance, as calculated from the fitted psychometric functions.

### Subjective quality description task

In each trial, a 2-second pulse train was delivered and participants were asked to indicate which subset of a list of 30 qualitative descriptors applied to the percept (word list shown in Figure 1d). These words were selected based on prior studies of language associated with natural and artificial somatosensation^14,56–62^. Participants were encouraged to select as many words as necessary to fully describe the sensation. Participants were instructed to use their own definitions for each descriptor, but to maintain consistency throughout each experimental session. For example, if two stimuli felt identical, they were asked to select the same subset of words for both trials. Stimulation was applied at one of 9 PFs and 3 PWs for each stimulus. Stimulation PF was set to 2, 5, 10, 20, 50, 100, 200, 500, and 1000 Hz, and PWs were selected in the same manner as in the frequency estimation task. Each stimulus was presented 20 times in pseudo-randomized order. A 2-second inter-trial interval was enforced. A “no sensation” option was provided for trials in which the participant did not feel the stimulus. Eleven electrode contacts were tested across all six participants. Stimulation conditions in which “no sensation” was selected in more than five trials were excluded from the analysis. We computed the proportion of times each descriptor word was selected for each stimulation condition for each contact after “no sensation” trials were removed. The result was a set of 30 proportions for each stimulation condition.

To compute within-participant consistency in word selection, we ran a correlation analysis of word proportions between the sets of contacts tested within the same participant. Those participants that only performed the subjective quality test with a single contact were excluded from this analysis. To calculate across-participant consistency in word selection, we first averaged the proportions across contacts for each participant, then computed the correlation in these proportions for each pairing of participants.

#### Dissimilarity matrices

Having observed that different participants used different words to describe the electrically-evoked sensations, we hypothesized that we could achieve a more generalizable representation of tactile quality by computing differences in descriptors between stimulation conditions. To reduce redundant or highly-correlated descriptors within each dataset, we ran a principal component analysis (PCA) on the word proportion vectors across all stimuli (a 30 x 27 data matrix) on a participant-by-participant basis and retained the components necessary to explain 95% of the variance. We then projected the stimuli onto this reduced dimensionality space and calculated the pairwise distances between them, resulting in a 27 x 27 distance matrix for each tested electrode contact (n=11) (Supplementary Figure 3a). To remove the influence of the number of dimensions on calculated distances, we normalized distances to the maximum inter-condition distance (which was set to 1). For some analyses (e.g. that shown in Figure 6a), we averaged the coordinates for each PF across PWs and then recomputed the distances, yielding a 9 x 9 distance matrix. For Supplementary Figure 3b, we averaged the coordinates for each PW across PFs and computed distances, yielding a 3 x 3 distance matrix.

To assess the within-participant consistency in dissimilarities, we computed the correlation between distance matrices obtained from different contacts for each participant. To assess across-participant consistency in dissimilarities, we first averaged the dissimilarities across contacts for each participant then computed the correlation in the dissimilarity matrices for each pair of participants.

To examine the quality dissimilarity between successive PF conditions, we computed the distance between the coordinates of two adjacent PFs in the reduced dimensionality space, then averaged these distances across PWs and contacts.

Finally, we performed a multivariate regression analysis with ΔPF, ΔPW, and their interaction as factors. We performed this analysis after averaging the 27 x 27 distance matrices across participants and contacts. The ΔPF and ΔPW factors were z-scored before running the regression to standardize the regression coefficients. This analysis was also performed with the data split into two sets: one for PFs below 50 Hz and one for PFs above 50 Hz. We verified that the regression model fit to the dissimilarity matrix averaged across contacts and participants yielded the same conclusions as one fit to data pooled across individual contacts and participants. We also fit regression models to individual participants’ dissimilarity matrices, and verified that the regression coefficients averaged across these models yielded the same conclusions (Supplementary Figure 4). We then performed a step-wise regression, in which we assessed whether PF accounted for a significant proportion of variance in the dissimilarity ratings after differences in perceived frequency had been regressed out. For this analysis, we first computed differences in perceived frequency for each stimulus pair from the across-contact averages of perceived frequency obtained in the first experiment. We then regressed dissimilarity against differences in perceived frequency ratings and regressed the residuals against z-scored ΔPF. This analysis was performed separately for stimuli below and above 50 Hz.

### Statistical Tests

#### Frequency estimations

We determined the influence of PW on the transition frequency between the rising and falling regimes of perceived frequency using a one-way ANOVA versus PW. We then split the data into two regimes: 0-50 Hz and above 50 Hz. For each regime, we ran a two-way ANOVA on the normalized frequency estimations pooled across contacts and participants with factors of PW and PF. All statistical analyses were performed in Minitab (Minitab, LLC; State College, PA, USA) with alpha level of 0.05.

#### Frequency discriminations

We studied the influence of PW on the JNDs and PSEs of the psychometric functions using repeated measures ANOVA with different PWs and different contacts as factors. JNDs and Weber fractions at 20Hz standard were compared to their counterparts at 50Hz using a two-sample t-test. All statistical analyses were performed in MATLAB with alpha level of 0.05.

#### Subjective quality reports

We ran a linear regression to determine how the number of words selected by each participant changed with PF. We compared within- and across-participant correlations in quality descriptors and dissimilarities using a one-way ANOVA with Tukey pairwise comparisons. We compared quality dissimilarities between successive PFs using a two-way ANOVA with Tukey pairwise comparisons with factors for PF and PW. These analyses, as well as the univariate and multivariate regressions, were performed in MATLAB and/or Minitab with alpha level of 0.05.

## Acknowledgements

We would like to thank the research participants for their time and dedication to advancing science and M. Schmitt for assistance with participant clinical care and study coordination. We would also like to thank C. Hughes, R. Gaunt, B. Delhaye, and T. Callier for their helpful feedback on an early version of the manuscript. This work was sponsored by the DARPA Biological Technologies Office (BTO) HAPTIX program under the auspices of Dr. A. Emondi through the Space and Naval Warfare Systems Center (Pacific contract no. NC66001-15-C-4041), by the U.S. Department of Veterans Affairs Rehabilitation Research and Development Service Program (Center #C3819C), and by the National Institute of Neurological Disorders and Stroke (5R01NS095251-05). The content is solely the responsibility of the authors and does not necessarily represent the official views of the listed funding institutions.

## Author Contributions

ELG designed the study, collected the data, analyzed the data, interpreted the results, and wrote the manuscript. BPC collected the data, analyzed the data, and wrote the manuscript. QH analyzed the data and wrote the manuscript. DJT advised on result interpretation and revised the manuscript. SJB designed the study, interpreted the results, and wrote the manuscript.

