## Supplemental information for "Frequency shapes the quality of tactile percepts evoked through electrical stimulation of the nerves"

### SUPPLEMENTARY INFORMATION

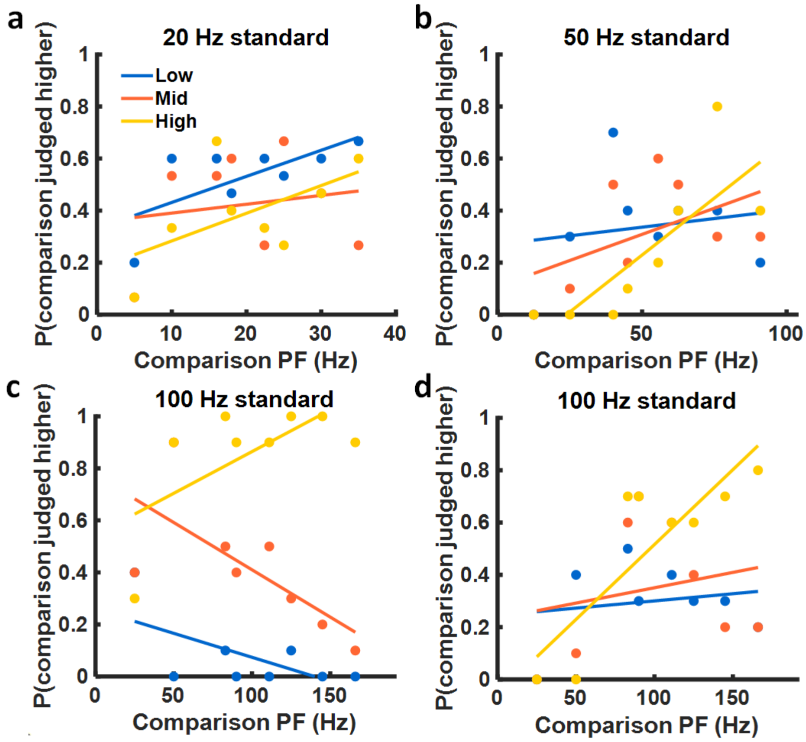


*Supplementary Figure 1. Examples of electrode contacts with poor frequency discrimination performance. Each panel depicts one contact in which the participant did not reach criterion performance. Lines are fitted to different comparison PWs (blue for low PW, orange for mid PW, and yellow for high PW). a) Contact G4 of participant LL02. b) Contact M2 of UL04. c) Contact M3 of UL04. d) Contact M5 of UL04.*

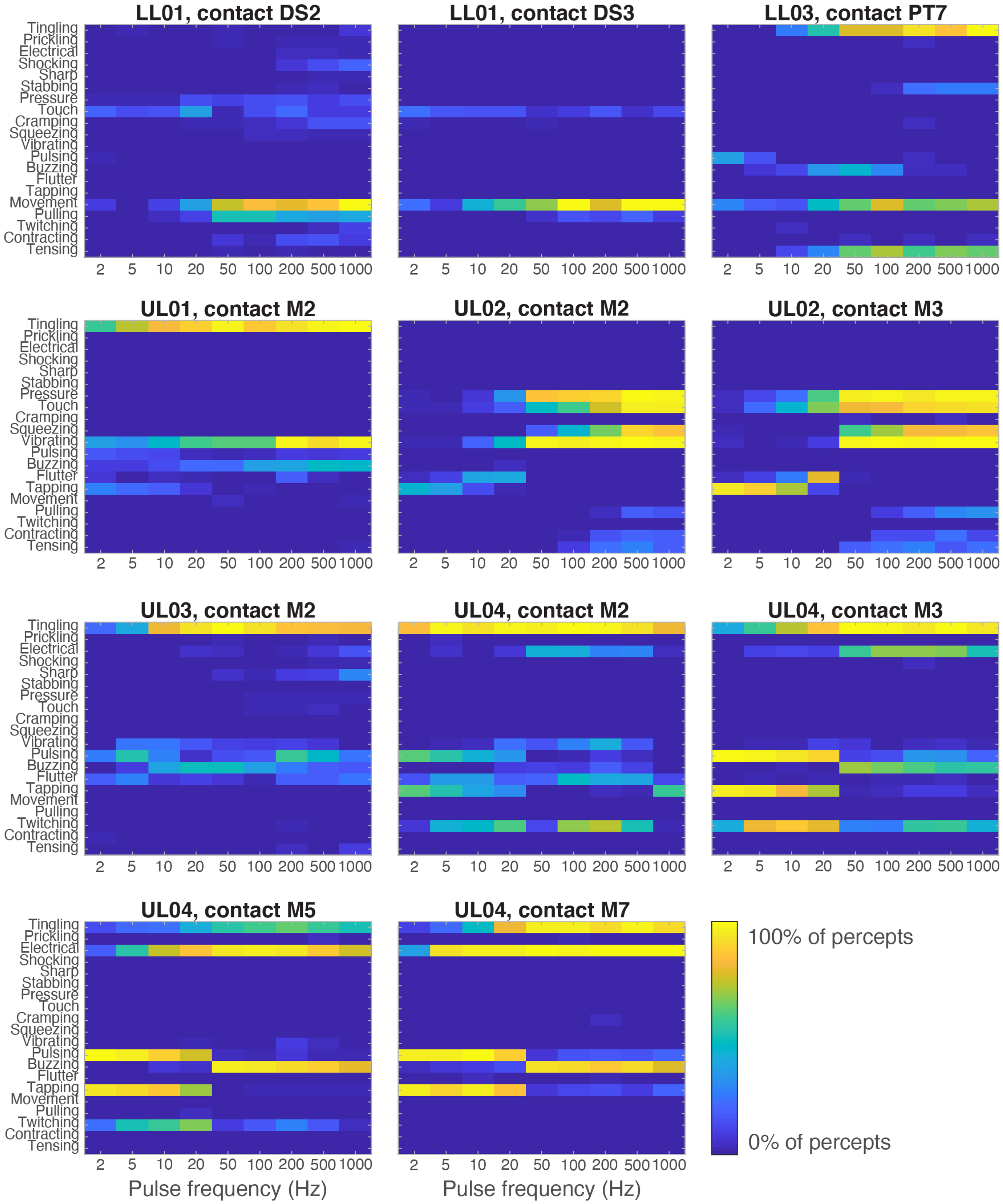


*Supplementary Figure 2. Effect of PF on qualitative reports for all participants and contacts (n=11). Each row corresponds to a descriptor. The color denotes the proportion of perceived trials at each stimulation PF in which the descriptor was reported.*

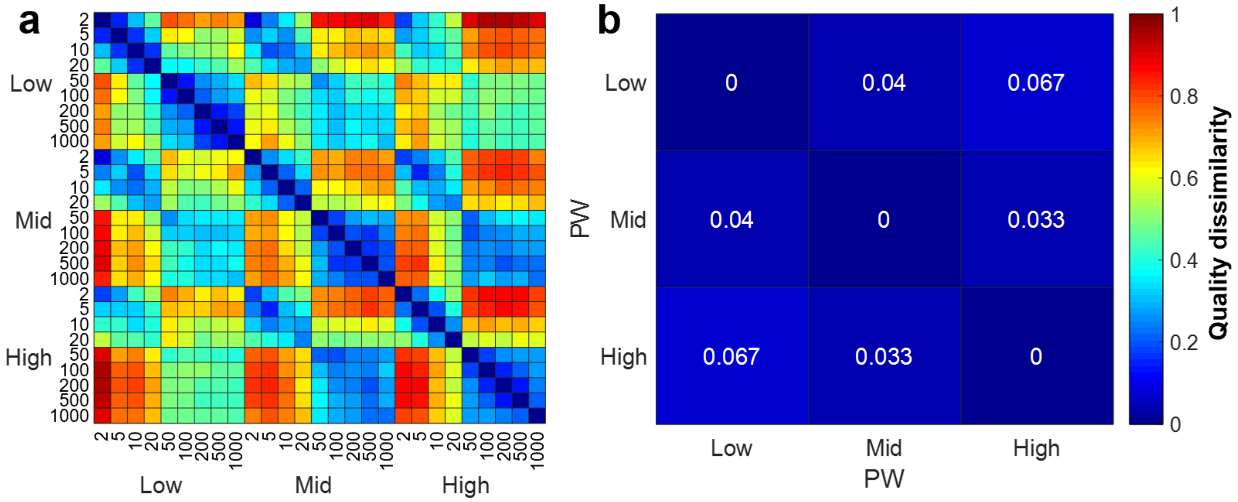


*Supplementary Figure 3. Perceived dissimilarities across stimulation conditions. a) Full 27x27 dissimilarity matrix, averaged across participants and contacts (n=11). Conditions are grouped first by PW (Low, Mid, High) and then by PF (in Hz). b) Perceived dissimilarity for all participants and contacts, averaged for each PW (Low, Mid, High) across PFs (n=11).*

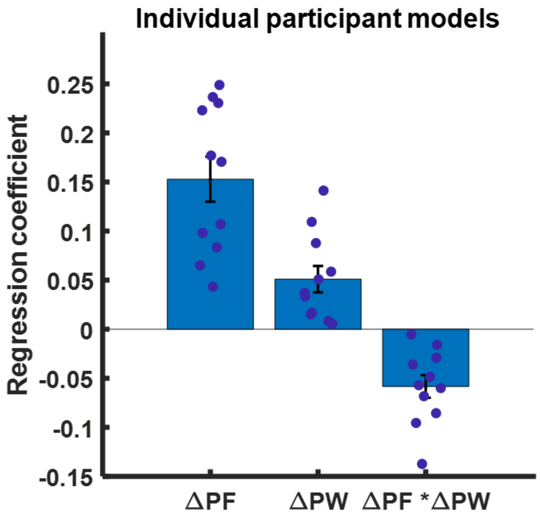


*Supplementary Figure 4. Regression coefficients for models fit to individual participants' dissimilarity data. Each regression model was a linear combination of ΔPF, ΔPW, and their interaction. The regression coefficients for individual models are denoted as dark blue filled circles, while the mean coefficients are shown as the light blue bars. Error bars denote the standard error of the mean (n=11).*

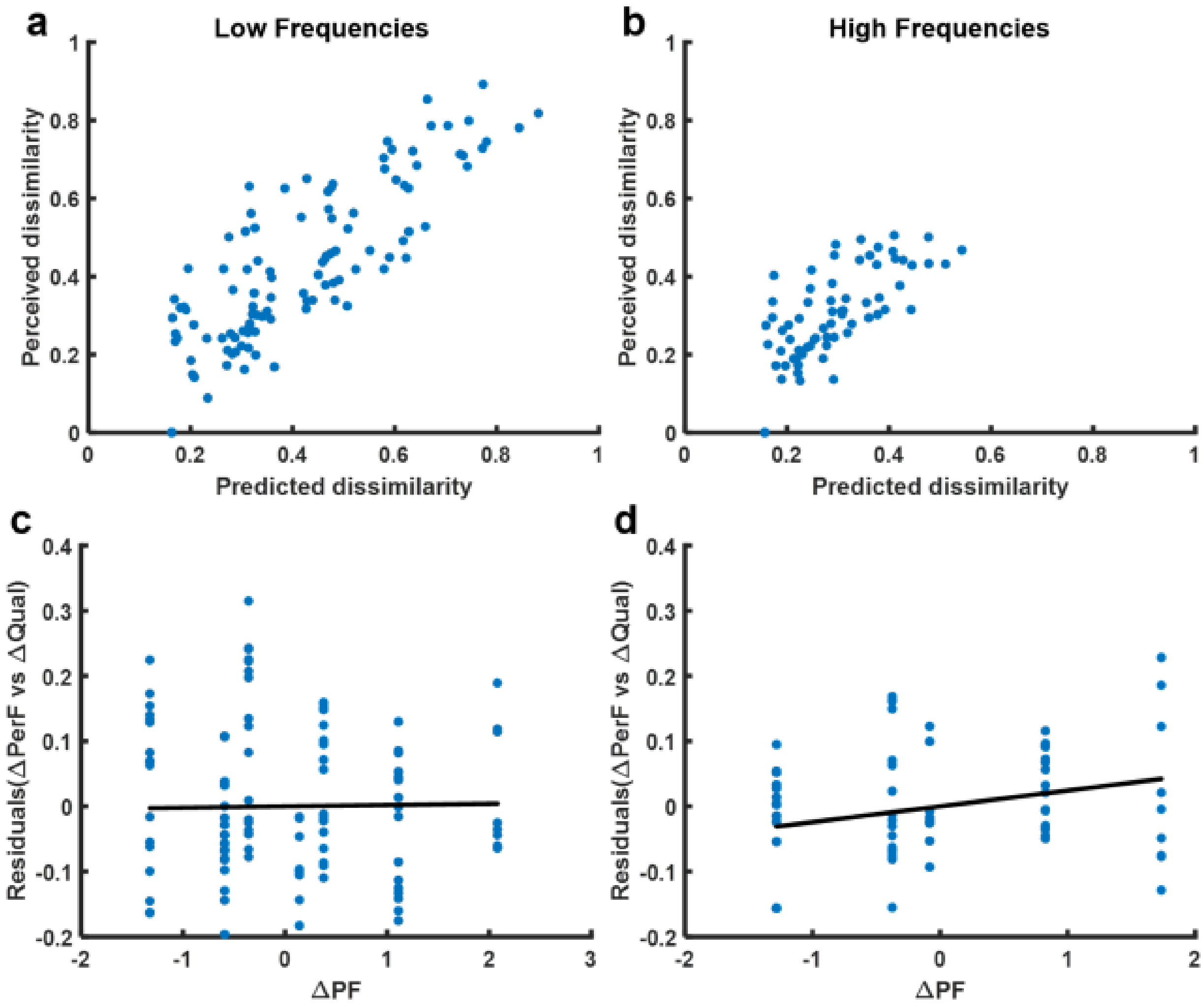


*Supplementary Figure 5. Effect of PF on perceived dissimilarity after removing the influence of perceived frequency. A univariate regression predicting quality dissimilarity based on perceived frequency was performed for low (a) and high (b) frequency stimuli. a,b) Perceived dissimilarity versus a linear prediction based on differences in perceived frequency. c,d) Residuals of the regression (shown in a and b) versus (z-scored) ΔPF. Solid black lines denote the regression of the residuals on to ΔPF. c) At low frequencies, the influence of ΔPF on dissimilarity was not significant after accounting for perceived frequency ($p = 0.8$). d) At high frequencies, ΔPF significantly impacted dissimilarity, even after accounting for perceived frequency ($p = 0.001$), but the effect was modest.*

*Supplementary Table 1. The selection rate for perceptual descriptors differed for low frequency and high frequency stimulation conditions. Selection rates are given as the percentage of total trials that each descriptor word was reported across all participants, contacts, and PWs. The top seven most frequently reported words for each frequency regime are shown. Note that since multiple words can be selected for a single trial, the selection rates will not sum to 100%.*

| Low frequency (<50 Hz) | | High frequency (≥50 Hz) | |
|---|---|---|---|
| **Descriptor** | **Selection Rate** | **Descriptor** | **Selection Rate** |
| Tapping | 36% | Tingling | 49% |
| Pulsing | 34% | Buzzing | 28% |
| Tingling | 29% | Electrical | 26% |
| Twitching | 15% | Vibrating | 22% |
| Electrical | 14% | Pressure | 13% |
| Flutter | 7% | Touch | 12% |
| Vibrating | 6% | Movement | 9% |

### SUPPLEMENTARY NOTES

We explored several possible mechanisms underlying the transition in quality perception ~50 Hz. One possibility is that the tactile nerve fibers cannot, as a population, phase lock to stimulation pulse trains beyond about 50 Hz. Indeed, the temporal patterning in the neural response likely drives the effect of stimulation PF on sensory quality. At high frequencies, stimulation pulses may encroach on the refractory period resulting from the neural response to the previous pulse. This phenomenon might then blur the temporal patterning in the population response. However, given that the refractory period is typically less than 5 ms[1] and that pulses are separated by 20 ms at 50 Hz, refractoriness is unlikely to play a critical role in the observed transition in quality.

Another possibility is that the spikes evoked in the afferent population by each pulse get desynchronized at proximal stages of the sensory pathway due to conduction delays, thereby blurring the temporal patterning at the population level. Indeed, neural conduction velocities vary across Aβ nerve fibers that mediate touch due to the natural variance in fiber diameters[2]. In an adult male, the resulting propagation delays for signals to travel from the fingertip to spinal cord range from 11 to 22 ms[3]. However, this jitter is not observed in the vibrotactile responses of neurons in the somatosensory cortex[4,5], which exhibit a high degree of phase locking, suggesting that some compensatory mechanism might eliminate this endogenous jitter along the way to the brain. The synchronized spikes delivered through a nerve cuff might then *become* desynchronized via this compensatory mechanism. Given that the intrinsic delay correction would span about 10 ms (to correct for delays ranging from 11 to 22 ms) and that the cuff electrode is positioned approximately halfway between the fingertip and spinal cord, the imposed jitter would span approximately 5 ms. We might then expect frequency discrimination to break down around 200 Hz. Thus, this conduction delay-mediated desynchronization mechanism is also unlikely to be solely responsible for the transition at 50 Hz.
